# The methylome and comparative transcriptome after high intensity sprint exercise in human skeletal muscle

**DOI:** 10.1101/2020.09.11.292805

**Authors:** Mohd Firdaus Maasar, Daniel C. Turner, Piotr P. Gorski, Robert A. Seaborne, Juliette A. Strauss, Sam O. Shepherd, Matt Cocks, Nicolas J. Pillon, Juleen R. Zierath, Andrew T. Hulton, Barry Drust, Adam P. Sharples

## Abstract

The methylome and transcriptome signature following exercise that is physiologically and metabolic relevant to sporting contexts such as team sports or health prescription scenarios (e.g. high intensity interval training/HIIT) has not been investigated. To explore this, we undertook two different sport/exercise relevant high-intensity sprint running protocols in humans using a repeated measures design of: 1) Change of direction (COD) versus; 2) straight line (ST) sprint exercise. We took skeletal muscle biopsies from the vastus lateralis 30 minutes and 24 hours post exercise followed by 850K methylation arrays and comparative analysis with recent sprint and acute aerobic exercise meta-analysis transcriptomes. Despite matched intensity (speed x distance and number of accelerations/decelerations) between COD and ST exercise, COD exercise elicited greater movement (GPS Playerload™), physiological (HR), metabolic (lactate) as well as central and peripheral (differential RPE) loading compared with ST exercise. The exercise response alone across both conditions evoked extensive alterations in the methylome immediately post and 24 hrs after exercise, particularly in MAPK, AMPK and axon guidance pathways. COD evoked a considerably greater hypomethylated signature across the genome compared with ST sprint exercise, particularly enriched in: Protein binding, MAPK, AMPK, insulin, and axon guidance pathways. A finding that was more prominent immediately post exercise. Comparative methylome analysis with sprint running transcriptomes identified considerable overlap, with 49% of the genes altered at the expression level also differentially methylated after COD exercise. After differential methylated region analysis, we discovered that VEGFA and its downstream nuclear transcription factor, NR4A1 had enriched hypomethylation within their promoter regions. VEGFA and NR4A1 were also significantly upregulated in the sprint transcriptome and meta-analysis of exercise transcriptomes. We confirmed increased mRNA expression of VEGFA, and considerably larger increases in the expression of canonical metabolic genes, PGC1-α and NR4A3, 3 hrs post COD vs. ST exercise. Overall, we demonstrate that increased physiological load via change of direction sprint exercise in human skeletal muscle evokes considerable epigenetic modifications that are associated with changes in expression of genes responsible for adaptation to exercise. These data imply that introducing changes in direction into high intensity running protocols could serve as an important modulator of a favourable epigenomic and transcriptomic landscape in response to exercise in athletes and trigger greater skeletal muscle remodelling through enhanced gene expression.

## Introduction

Epigenetic modifications, particularly DNA methylation, is associated closely with altering gene expression in response to exercise in human skeletal muscle^1^. Genome-wide DNA methylation analysis after acute resistance exercise, training, detraining and retraining in humans has demonstrated that epigenetic modifications can be retained and therefore skeletal muscle possesses an epigenetic memory^1^. Resistance exercise preferentially reduces (hypomethylates) the DNA methylome^2, 3^ and increases gene expression across the transcriptome^1^. With even greater hypomethylation observed if exercise training had been undertaken previously, such as following retraining^2, 3^. Candidate genes demonstrating this epigenetic profile, such as UBR5 are important regulators of skeletal muscle mass^4, 5^. Therefore, alterations in DNA methylation signatures are emerging as important drivers for the molecular response to exercise and subsequent physiological adaptation.

Methylation of DNA occurs most commonly in cytosine-guanine (C-G) base pairings called CpG sites. CpG islands are CpG rich locations that occur more frequently in gene regulatory regions, such as promoter or enhancer regions. In humans the majority of CpG sites are methylated (70-80%)^6, 7^. When methylation is present or increases (hypermethylation), particularly in promoters or enhancers, this prevents transcription factors from binding to these portions of the gene to enable gene expression^8^. In contrast, loss of methyl groups (hypomethylation) from these regions can enable/increase gene expression.

Aerobic exercise training alters the skeletal muscle methylome in humans^9^ and preferentially hypomethylates the genome in type-II diabetics^10^ and obese populations^11^. At the targeted gene level, acute aerobic exercise, particularly at higher intensities (80% vs. 40% VO_2_ max) hypomethylates PGC-1α, a well-established regulator of mitochondrial biogenesis. With the same trend observed for associated metabolic pathway genes; TFAM, PDK4 and PPAR-δ^12^, together with a corresponding increase in their gene expression^12^. However, there is limited analysis of DNA methylation across the genome after acute aerobic exercise in healthy human skeletal muscle. Furthermore, there is no temporal assessment of the methylome post-acute aerobic exercise, such as after 24 hrs or after alterations in exercise intensity. Finally, the methylome signature following exercise that is physiologically and metabolic relevant to sporting contexts such as team sports or health prescription scenarios (e.g. high intensity interval training/HIIT) in the general public has not been investigated.

In the present study, we devised a repeated measures sprint shuttle running protocol in humans that was physiologically relevant to exercise movements within many sporting contexts and health prescription settings. In the design there were two different sprint shuttle trials: 1) Change of direction (COD) and, 2) straight line (ST) sprint exercise that were both matched for classically defined intensity measures (speed x distance) and the number of acceleration and deceleration movements. However, despite this matched intensity between trials, in the COD exercise protocol greater movement (GPS Playerload™), physiological (HR), metabolic (lactate), subjective central and peripheral (differential RPE) loading was elicited compared with ST sprint running. Therefore, the comparison of COD versus ST exercise represented a suitable model to investigate the time-course of the DNA methylome following increased loading exercise in human skeletal in a physiologically relevant model. Therefore, we first aimed to investigate differential DNA methylation across the genome between these two trials with dedicated genome-wide analyses for enriched gene ontologies, pathways and chromosomal regions 30 minutes and 24 hrs after COD versus ST exercise. Furthermore, we undertook comparative analysis with recently published transcriptome data sets after acute sprint exercise in humans^13^, and also with recent meta-analysis that combines transcriptome data from 66 published data sets in the most comprehensive exercise transcriptome profiling to date^14^. Subsequently, the secondary aim was to determine if the alterations across the methylome between COD versus ST exercise were also seen at the transcriptome level. With the ultimate aim of identifying epigenetic regulators in the response to acute exercise at the pathway and candidate gene level. Overall, the present study demonstrated that increased physiological load, incorporating changes of direction into sprint shuttle exercise in human skeletal muscle, evoked an extensive epigenetic signature that was associated hypomethylation and with changes of expression of gene pathways responsible for the adaptation to exercise (AMPK / MAPK / insulin/ axon guidance / protein binding pathways), as well as genes at the candidate level (VEGFA, NR4A1, NR4A3 and PGC1-α). This study therefore suggests that introducing changes of direction into running protocols could serve as an important modulator in promoting a favourable epigenomic and transcriptomic response to acute exercise. Finally, an applied implication of the study suggests that by incorporating changes of direction into exercise regimes overtime (COD training) may ultimately improve performance within a sporting context or health outcomes of populations with dysfunction in the identified (MAPK, MAPK, insulin) gene pathways.

## Methods

### Experimental overview

Following preliminary visits to the laboratory to complete screening and pre-test assessments/familiarisation a repeated measures design was used for all participants to complete two experimental trials; a 180-degree change of direction (COD) performed over a distance of 5 m to volitional exhaustion and a repeated straight line 5 m sprint protocol (ST) performed at the same intensity (running speed) until an equivalent distance to that covered in COD was completed (speed: 3.2 ± 0.1m.s^-1^; distance: 594 ± 35 m). The individual responses to these exercise bouts were described from a movement (GPS player load), physiological (HR), metabolic (lactate) and subjective perspective (central and peripheral fatigue / rating of perceived exertion -RPE) to holistically characterize the ‘load’ demand associated with the activity. Muscle biopsies were obtained from the lateral portion of the vastus lateralis muscle for genome wide DNA methylation analysis, comparative transcriptome and targeted gene expression, described in detailed below.

### Participants

A group of five healthy well-trained, male team sports players (age: 26 ± 2 yrs; height: 1.7 ± 0.1 m; weight: 76.3 ± 11.5 kg) were recruited to participate in this study. After giving informed consent, all participants were screened using physical activity and readiness to exercise questionnaire and a pre-biopsy medical assessment checklist. The information was used by the research team and a qualified medical practitioner to confirm inclusion criteria and eligibility for the current study. All participants included were free from any musculoskeletal injuries for the last six months. Participants were required to refrain from strenuous exercise 24 hours prior to testing, alcohol for 48 hours prior to exercise and caffeine for 12 hours prior to exercise. Ethical approval was granted from the institutional ethics review board/committee for the study (ref. 19/SPS/028).

### Pre testing assessments/familiarisation

#### Max Running Speed Test

All participants attended the laboratory, a maximum of two occasions, to complete a pre-test assessment to determine the running speed to be used in the experimental protocols and to complete familiarisation with the relevant measurement tools. Each participant was required to perform a 6 x 5m shuttle run with a 180-degree change of direction at their self-chosen maximal speed. These runs were performed until the speed at which the shuttles were completed demonstrated a coefficient of variation of 5% or lower. This intensity was also re-evaluated on a second visit to ensure that participants could re-produce the required running speed identified in the first session to the same accuracy (3.21 ± 0.1 m.s^-1^). This speed was then used for the completion of all subsequent shuttle runs, programmed into a computer derived audible beep. All participants also performed 10 repetitions in a preliminary trial of COD with 30-second rest in between repetitions. This acted as familiarisation to both the movement patterns required for the protocol and the approach to be used to evaluate the subjective responses to exercise.

### Experimental exercise protocols

Participants performed two experimental protocols following an overnight fast; COD and ST on separate occasions separated by a minimum of 2 weeks. All participants completed the COD protocol first followed by ST. Both protocols were preceded by the completion of a standardised 10 min warm-up which consisted of 5 min slow running to elevate HR. This activity was followed by dynamic and static stretching to increase range of movement and readiness for the specific movement pattern of the running protocol. The warmup was led by the researcher. The intensity (running speed) of COD was controlled using a computer derived audible beep determined based on the maximum individual speed collected during the pre-test assessment. After each repetition (6 x 5 m runs separated by a 180° COD), a 30s rest period was completed. COD was performed until voluntary exhaustion (defined as the point at which the participant could no longer complete the required shuttle distance at the pre-determined speed). The ST sprints were performed at an equivalent intensity to that used for COD. ST sprints required individuals to perform repeated 5 m sprints in a forward direction over a 30 m course. To maintain a similar acceleration and deceleration profile to that of the COD participants were required to come to a stop and then immediately set off again after each 5 m sprint. Computer derived audible beeps were again used to “pace” the completion of the bouts of activity. Participants maintained ST sprinting until the volume of activity matched that completed at voluntary exhaustion in the COD trial.

Dependent variables associated with movement, physiological and subjective outcomes were recorded to comprehensively describe the exercise in COD and ST and its associated demand. Prior to the exercise, an athlete tracking unit that included a micro-electrical mechanical device and global positioning system (Catapult S5, Australia) was fitted into a purpose made vest and placed on each participant so it was positioned between the shoulder blades. All participants used the same tracking unit throughout the testing session to avoid inter-unit variability of the measurement^15^. A short-range radio telemetry device to monitor heart rate was also fitted (Polar T31 Kempele, Finland) according to manufactures guidelines. Both movement and heart rate were recorded continuously throughout the exercise session. Following the completion of the experimental protocol, data was downloaded using manufacturers software (Catapult Openfield software pack) to obtain the peak heart rate observed in COD and ST trials and the peak Playerload™ (the instantaneous rate of change of acceleration divided by a scaling factor, used as an indicator of the overall movement demand^15^). During both experimental protocols between the 30s rest periods between sprints, participants were asked to self-rate their differential RPE for both local (legs) and central (breathlessness) exertion using a Centimax 100 scale^16^. This data provided a subjective evaluation of the central and peripheral demand associated with the trials. The peak differential RPE for both legs and breathlessness were used to compare subjective ratings between COD and ST. Capillary blood lactate was determined at rest (baseline) and post exercise from a fingertip blood sample for each participant. Following a puncture made using a lancet (Accu-Chek, Safe-T Pro Plus) a blood sample was drawn from the fingertip, using 20 μl blood capillary tubes. The blood sample was immediately placed and mixed into the lactate haemolysing solution cup, to prevent blood sample from clotting. All samples were analysed using Biosen C-Line “EKF diagnostic” device. Pre-blood lactate sample was drawn at rest before the warm-up, and post-blood lactate was drawn 5 minute following the completion of the exercise trial.

### Skeletal muscle biopsies

Muscle biopsies were taken from the lateral portion of the vastus lateralis muscle under local anaesthetic with Marcain using the conchotome technique. Due to ethical considerations of multiple biopsies from the same leg over a short time period, within each trial muscle biopsies were taken from both legs (detailed below). Upon arrival at the laboratory under overnight fasted conditions, the pre-exercise muscle biopsy was collected from the non-dominant leg. Post-exercise muscle biopsies were taken (from the dominant leg) 30 minutes following the completion of COD and ST exercise. A final biopsy was taken 24-hour post-exercise again following an overnight fast (from the non-dominant leg). EMG analysis confirmed that there were no significant differences in muscle activation between dominant and non-dominant leg during COD exercise.

### Tissue Homogenization, DNA isolation and bisulfite conversion

Due to cost, tissue samples from three out of the five well-trained male team sports players (age: 27 ± 2 yrs; height: 1.7 ± 0.1 m; weight: 73.5 ± 11.1 kg) were randomly taken forward for DNA methylome analysis at baseline and immediately post and 24 hrs post in both COD and ST trials. Tissue samples were homogenized for 45 seconds at 6,000 rpm × 3 (5 minutes on ice in between intervals) in lysis buffer (180 μl buffer ATL with 20 μl proteinase K) provided in the DNeasy spin column kit (Qiagen, UK) using a Roche Magnalyser instrument and homogenization tubes containing ceramic beads (Roche, UK). Cells were lysed in 180 μl PBS containing 20 μl proteinase K. The DNA was then bisulfite converted using the EZ DNA Methylation Kit (Zymo Research, CA, United States) as per manufacturer’s instructions.

### Infinium MethylationEPIC BeadChip Array

All DNA methylation experiments were performed in accordance with Illumina manufacturer instructions for the Infinium Methylation EPIC BeadChip Array. Methods for the amplification, fragmentation, precipitation and resuspension of amplified DNA, hybridisation to EPIC beadchip, extension and staining of the bisulfite converted DNA (BCD) can be found in detail in our open access methods paper^3^. EPIC BeadChips were imaged using the Illumina iScan System (Illumina, United States).

### DNA methylation analysis, CpG enrichment analysis (GO and KEGG pathways), differentially modified region analysis and Self Organising Map (SOM) profiling

Following DNA methylation quantification via MethylationEPIC BeadChip array, raw .IDAT files were processed using Partek Genomics Suite V.7 (Partek Inc. Missouri, USA) and annotated using the MethylationEPIC_v-1-0_B4 manifest file. We first checked the average detection p-values. The mean detection p-value for all samples was 0.000141, and the highest was 0.00024 (Suppl. Figure **1a**), which is well below the recommended 0.01 in the Oshlack workflow^17^. We also examined the raw intensities/signals of the probes, that demonstrated an average median methylated and unmethylated signal of over 11.5 (11.74) and the difference between the average median methylated and average median unmethylated signal was 0.28, well below the recommended difference of less than 0.5^17^. Upon import of the data into Partek Genomics Suite we filtered out probes located in known single-nucleotide polymorphisms (SNPs) and any known cross-reactive probes using previously defined SNP and cross-reactive probe lists identified in earlier EPIC BeadChip 850K validation studies^18^. Although the average detection p-value for each sample across all probes was on average very low (no higher than 0.00024) we also excluded any individual probes with a detection p-value that was above 0.01 as recommended previously^17^. Therefore, out of a total of 865,860 probes, removal of known SNPs, cross-reactive probes and those with a detection p-value above 0.01 resulted in a final list of 809,877 probes analysed. Following this, background normalisation was performed via functional normalisation (with noob background correction), as previously described^19^. Following functional normalisation, we also undertook quality control procedures via principle component analysis (PCA), density plots by lines as well as box and whisker plots of the normalised data for all samples (Suppl. Figures **1b, c, d** respectively). We confirmed that no samples demonstrated large variation (variation defined as any sample above 2 standard deviations (SDs) – depicted by ellipsoids in the PCA plots (Suppl. Figure **1b**) and/or demonstrating any differential distribution to other samples, depicted in the signal frequency by lines plots (Suppl. Figure **1c**)). Therefore, no outlier samples were detected. We used participants baseline samples from the COD trial only for array analysis, as these were resting biopsies from the same participant under the same conditions before each trial. We also confirmed that baseline samples were not different between COD and ST trials by running a participants COD and ST trial baseline samples and demonstrating that the PCA showed very little variation between the baseline samples taken before the COD trial and the baseline sample taken before the ST trial (Suppl. Figure **1c**-Samples highlighted with an arrow). Following normalisation and quality control procedures, we undertook differentially methylated position (DMP) analysis by converting β-values to M-values (M-value = log2(β / (1 - β)), as M-values show distributions that are more statistically valid for the differential analysis of methylation levels^20^. We performed a 2-way ANOVA for ‘Trial’ (ST vs. COD) x time (baseline, post, 24 h) with planned contrasts of: 1) COD Post vs. ST post, 2) COD 24 h vs ST 24 h). We also explored the ANOVA main effects for both ‘Trial’ to investigate the effect of trial alone (effect of COD vs. ST across time), and ‘Time’ (baseline, post and 24 h) with planned contrasts (post vs. baseline, baseline vs 24 h, post vs. 24 h) to investigate the effect of time alone (exercise effect alone over time in both COD and ST groups). Any differentially methylated CpG position (DMP) with an adjusted P value of ≤ 0.01 was deemed significant. We then undertook CpG enrichment analysis on these differentially methylated CpG lists within gene ontology (GO) and KEGG pathways^21, 22, 23^ using Partek Genomics Suite, Partek Pathway and Revigo (tree-maps for GO terms). Differentially methylated region (DMR) analysis, that identifies where several CpGs are consistently differentially methylated within a short chromosomal location/region, was undertaken using the Bioconductor package DMRcate (DOI: <u>10.18129/B9.bioc.DMRcate</u>). Finally, in order to plot/visualise temporal changes in methylation across the time-course post exercise (COD vs. ST) we implemented Self Organising Map (SOM) profiling of the change in mean methylation within each condition using Partek Genomics Suite.

### Comparative transcriptome analysis

We overlapped the significant differential sprint methylome data from the present study with significantly expressed sprint transcriptomes^13^ and also with a meta-analysis of acute aerobic exercise transcriptomes^14^. Venn diagram analysis was used to generate gene lists of overlap between differential methylome and transcriptome data. We also used this approach to confirm the genes that were identified to be hypomethylated in the present study and upregulated in the sprint transcriptome^13^, that were also upregulated in the aerobic exercise transcriptome meta-analyses^14^.

As well as genes that were hypermethylated in the present study and down-regulated in the transcriptome data sets. Metamex.eu^14^ was used to confirm the genes logfold change and significance (with an FDR of p ≤ 0.05) in the meta-analysis transcriptome data. In particular, we selected the criteria of healthy skeletal muscle up to 24 hrs post exercise in order to match to the design of the present study. To generate a list of genes that were significantly differentially methylated in the present study and significantly regulated at the expression level in transcriptome meta-analysis after acute aerobic exercise (but not identified in the sprint transcriptome)^13^ we conducted new analysis of acute aerobic exercise transcriptome datasets from the meta-analysis^14^. Therefore we also analysed full significant genes lists (adj. p value of ≤ 0.05) that were up- or down-regulated in transcriptome meta-analysis after acute aerobic exercise and overlapped these lists with our most significant methylated DMP analysis. New correlation analysis was performed on the non-publicly available acute aerobic meta-analysis data^14^ for the genes identified in the present manuscript at the methylome, comparative transcriptome and targeted gene level (RT-PCR below) to be significantly altered (including VEGFA, NR4A1, NR4A3 and PGC1-α).

### RNA isolation, primer design & gene expression analysis

Skeletal muscle tissue muscle from the same samples as the DNA methylome analysis across both trials (ST and COD) at baseline, 3h and 24 h post was homogenised in tubes containing ceramic beads (MagNA Lyser Green Beads, Roche, Germany) and 1 ml Tri-Reagent (Invitrogen, Loughborough, UK) for 45 seconds at 6,000 rpm × 3 (and placed on ice for 5 minutes at the end of each 45 second homogenization) using a Roche Magnalyser instrument (Roche, Germany). RNA was then isolated as per Invitrogen’s manufacturer’s instructions for Tri-reagent. Then a one-step RT-PCR reaction (reverse transcription and PCR) was performed using QuantiFast SYBR Green RT-PCR one-step kits on a Rotorgene 3000Q. Each reaction was setup as follows; 4.75 μl experimental sample (7.36 ng/μl totalling 35 ng per reaction), 0.075 μl of both forward and reverse primer of the gene of interest (100 μM stock suspension), 0.1 μl of QuantiFast RT Mix (Qiagen, Manchester, UK) and 5 μl of QuantiFast SYBR Green RT-PCR Master Mix (Qiagen, Manchester, UK). Reverse transcription was initiated with a hold at 50°C for 10 minutes (cDNA synthesis) a nd a 5-minute hold at 95°C (transcriptase inactivation and initial denaturation), before 40-50 PCR cycles of; 95°C for 10 sec (denaturation) followed by 60°C for 30 sec (annealing and extension). Primer sequences for genes of interest and reference genes were. VEGFA Fwd ACGGTCCCTCTTGGAATTGG, Rvse CTAATCTTCCGGGCTCGGTG; NR4A1 Fwd GGTGACCCCACGATTTGTCT, Rvse GGCTTATTTACAGCACGGCG; NR4A3 Fwd GACGTCGAAACCGATGTCAG, Rvse TTTGGAAGGCAGACGACCTC, PGC1-α Fwd TGCTAAACGACTCCGAGAA, Rvse TGCAAAGTTCCCTCTCTGCT; RPL13a Fwd GGCTAAACAGGTACTGCTGGG, Rvse AGGAAAGCCAGGTACTTCAACTT. All primers were designed to yield products that included the majority of transcript variants for each gene as an impression of total changes in the gene of interests’ expression levels. All genes demonstrated no unintended gene targets via BLAST search and yielded a single peak after melt curve analysis conducted after the PCR step above. All relative gene expression was quantified using the comparative Ct (^ΔΔ^Ct) method^24^. The baseline sample for each participant and their own reference gene sample was used as the calibrator conditions. The average, standard deviation and variations in Ct value for the RPL13a reference gene demonstrated low variation across all samples (mean ± SD, 21.04 ± 1.49, 7.1% variation) for the analysis. The average PCR efficiencies for VEGFA, NR4A1, NR4A3 and PGC1-α were comparable (90 ± 5.2, 89.2 ± 4.2, 91.3 ± 6.4%, 92.5 ± 1.9 variation) with the reference gene RPL13a (89.1 ± 4.8%). Statistical analysis genes were performed using a 2-way ANOVA with fisher post-hoc comparisons at the level of P value of ≤ 0.05 using Graphpad.

## Results

### Change of direction exercise elicits increased loading compared with ST exercise

Change of direction exercise increased mean movement loading (GPS Playerload™) (3.35 ± 0.07 vs. 2.93 ± 0.08 au; p ≤ 0.001), physiological loading (heart rate post exercise) (159 ± 16.3 vs. 137 ± 13.5 bpm; *p* ≤ 0.001), metabolic loading (lactate 5 minutes post exercise) (8.45 ± 1.7 vs. 1.69 ± 0.68 mmol/L p ≤ 0.05), subjective (differential RPE) peripheral loading (47 ± 24 vs. 10 ± 6; *p* ≤ 0.001) and central loading (62 ± 29 vs. 10 ± 6; *p* ≤ 0.001) versus straight line exercise.

### Exercise alone evokes a hypermethylated response immediately post exercise followed by hypomethylated signature by 24 h, particularly in MAPK, AMPK and Axon Guidance pathways

A 1-way ANOVA for ‘time’ (baseline, immediately post, 24 h after exercise), that analyses the main effect of sprint exercise alone over time across both experimental trials (COD and ST trials) identified a list of **7,612** significantly differentially methylated CpG positions (DMP’s) (**Suppl. File 1a**). Within this ‘exercise alone’ list, 1,076 DMPs were promoter associated with 1,651 located in CpG islands. Planned contrasts within this ANOVA for baseline vs. post exercise time points, also identified **4,952** significant DMPs (Suppl. **File 1b**). Where a larger number of DMPs were hypermethylated post exercise versus baseline, with 3287 hyper- and 1665 hypo-methylated. At 24 hrs (24 h vs. baseline contrast) there was **6,638 DMPs** (**Suppl. File 1c**), that demonstrated a shift towards a hypomethylated profile at 24 h, with 3,617 hypermethylated vs. 3,021 hypomethylated DMPs. Furthermore, this shift was confirmed in the 24h vs. post contrast, that identified a DMP list of **13,139 CpGs** (**Suppl. File 1d**), with a predominately hypomethylated profile, of 8,413 hypomethylated vs. 4726 hypermethylated DMPs. These data therefore suggest that immediately post exercise (across both experimental trials, COD and ST) DMPs favoured hypermethylation, however, after that point and up to 24 hrs there was shift towards a hypomethylated profile. This observation can be visualised as a hierarchical heat map in **Figure 1a**. Finally, SOM temporal analysis of the main effect for ‘time’ **7,612 DMP**s also confirmed the above analysis and suggested that the largest group of DMPs (e.g. 3,052 CpG’s highlighted in red and located in SOM profile; **Figure 1b**) were hypermethylated immediately post exercise, with the same DMPs demonstrated a shift to a hypomethylated profile 24 h after exercise.

**Figure 1.**
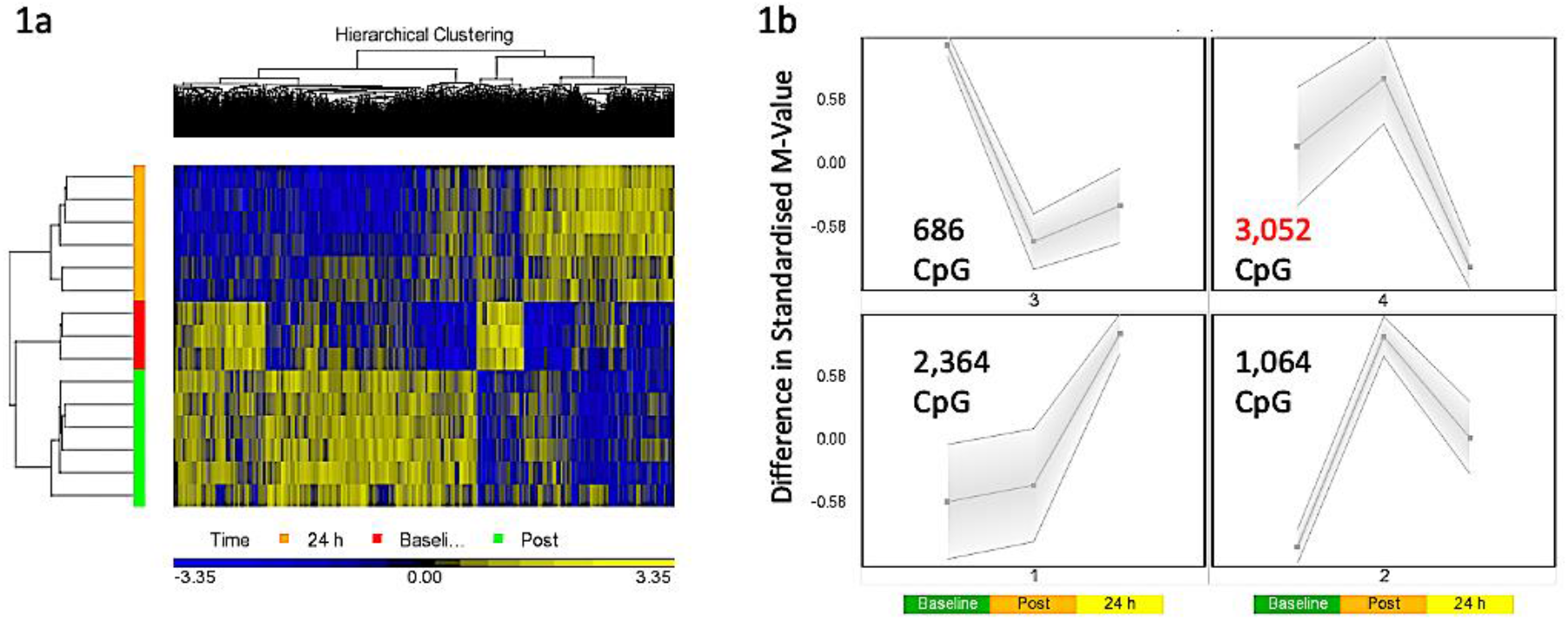
**a.** Hierarchical heat map for analysis of DMPs over exercise time in both trials (Red – Baseline, Post-Green, 24 h – Orange). **b.** SOM temporal analysis of DMPs over exercise time in both trials.

We next wished to identify which gene ontology (GO) terms and KEGG pathways were significantly enriched over the time-course of baseline, post and 24 following exercise (across both experimental trials (COD and ST). Indeed, GO enrichment analysis of the post vs. baseline contrast (**Suppl. File 1e**) identified that there was significant hypermethylation enrichment of DMPs in top 5 GO terms: 1) Neuron Part, 2) developmental process, 3) cellular component organisation, 4) regulation of neuron projection development and, 5) cellular component organisation or biogenesis (**Suppl. File 1f, g, h, i, j** respectively). For the 24 h versus baseline contrast (**Suppl. File 1k**) DMP hyper/hypomethylation was more balanced in top GO 5 terms: 1) Protein binding, 2) binding, 3) organelle, 4) intracellular part, and 5) intracellular organelle **(Suppl. File 1l, m, n, o, p).** Then at 24 h compared with post exercise (**Suppl. File 1q**) top 5 GO terms enriched predominantly hypomethylated were: 1) Developmental process, 2) anatomical structure development, 3) anatomical structure morphogenesis, 4) regulation of signaling and, 5) regulation of cell communication (**Suppl. File 1 r, s, t, u, v**).

For KEGG pathway analysis, in post vs. baseline contrasts (**Suppl. File 2a**) there was significant enrichment of hypermethylation of DMPs in top 5 KEGG pathways of: 1) MAPK signaling pathway, 2) axon guidance, 3) human papillomarvirus infection, 4) small cell lung cancer and, 5) insulin secretion (**Suppl. Files 2b, c, d, e, f**). At 24 h vs. baseline contrast (**Suppl. Files 2g**) top 5 enriched KEGG pathways were: 1) MAPK signaling pathway, 2) AMPK signaling pathway, 3) lysine degradation, 4) phosphonate and phosphate metabolism and, 5) aldosterone synthesis and secretion (**Suppl. Files, 2h, i, j, k, l**). At 24 h vs. post exercise (**Suppl. Files 2m**) hypomethylated profile enriched in top 5 KEGG pathways: 1) axon guidance, 2) cGMP-PKG signaling pathway, 3) focal adhesion, 4) rap1 signaling, and 5) adrenergic signaling in cardiomyocytes’ (**Suppl. Files, 2n, o, p, q, r**).

Finally, DMR analysis of the post vs. baseline (**Suppl. File 2s**), 24 h vs. baseline (**Suppl. File 2t**) and 24 h vs. post contrasts (**Suppl. File 2u**) identified several regions located in or close to annotated genes with enriched (multiple CpGs) differential methylation. Including enriched hypermethylation of genes: RBMXL1, ALDH3A1, EVA1A, MiR3928, RNF185, ANAPC10 and ABCE1 (post vs. baseline) and c12orf42, HSPD1, HSPE1, SALL1, CCNDN2 (24 h vs. baseline) and enriched hypomethylation in genes: ZIC1, ZIC4, TBX15, NAV2 and OXT (24h vs. post).

### Change of direction exercise evokes a greater hypomethylated signature in protein binding and axon guidance pathways compared with straight line exercise

Analysing the main effect for ‘Trial’ (COD vs. ST sprint exercise across all time points) identified **13,218** significant DMPs (**Suppl. File 3a**) favouring hypomethylation over hypermethylation (8,627 hypo-vs 4654 hyper-methylated). Undertaking hierarchical clustering of these DMPs enabled visualisation of the predominance in hypomethylation in COD versus the ST sprint exercise trials (**Figure 2a**). A large number of DMPs 4,221 (out of 13,218) were promoter associated, with 5,304 (out of 13,218) located in CpG islands. Undertaking gene ontology (GO) pathway enrichment within this **13,218 DMP** list (**Suppl. File 3b**). The top 5 GO terms enriched for hypomethylation in COD vs. ST sprint exercise were: 1) intracellular part, 2) protein binding, 3) binding, 4) organelle and, 5) intracellular organelle part (**Suppl. File 3c, d, e, f, g**), predominantly within overarching ‘cellular component’ (**Figure 2b**) and ‘molecular function’ gene ontologies (**Figure 2c**). The top KEGG pathways **(Suppl. File 3h)** predominantly hypomethylated were: 1) Axon guidance (**Figure 2d**), 2) cell cycle, 3) endocytosis, 4) pathways in cancer and, 5) spliceosome (**Suppl. Files 3 i, j, k, l, m** respectively). DMR analysis also identified that the genes: E2F3, BRD2, MUS81 and CFL1 had enriched hypomethylation of multiple DMPs in short chromosomal regions for these genes in COD versus ST sprint exercise (**Suppl. File 3n**).

**Figure 2.**
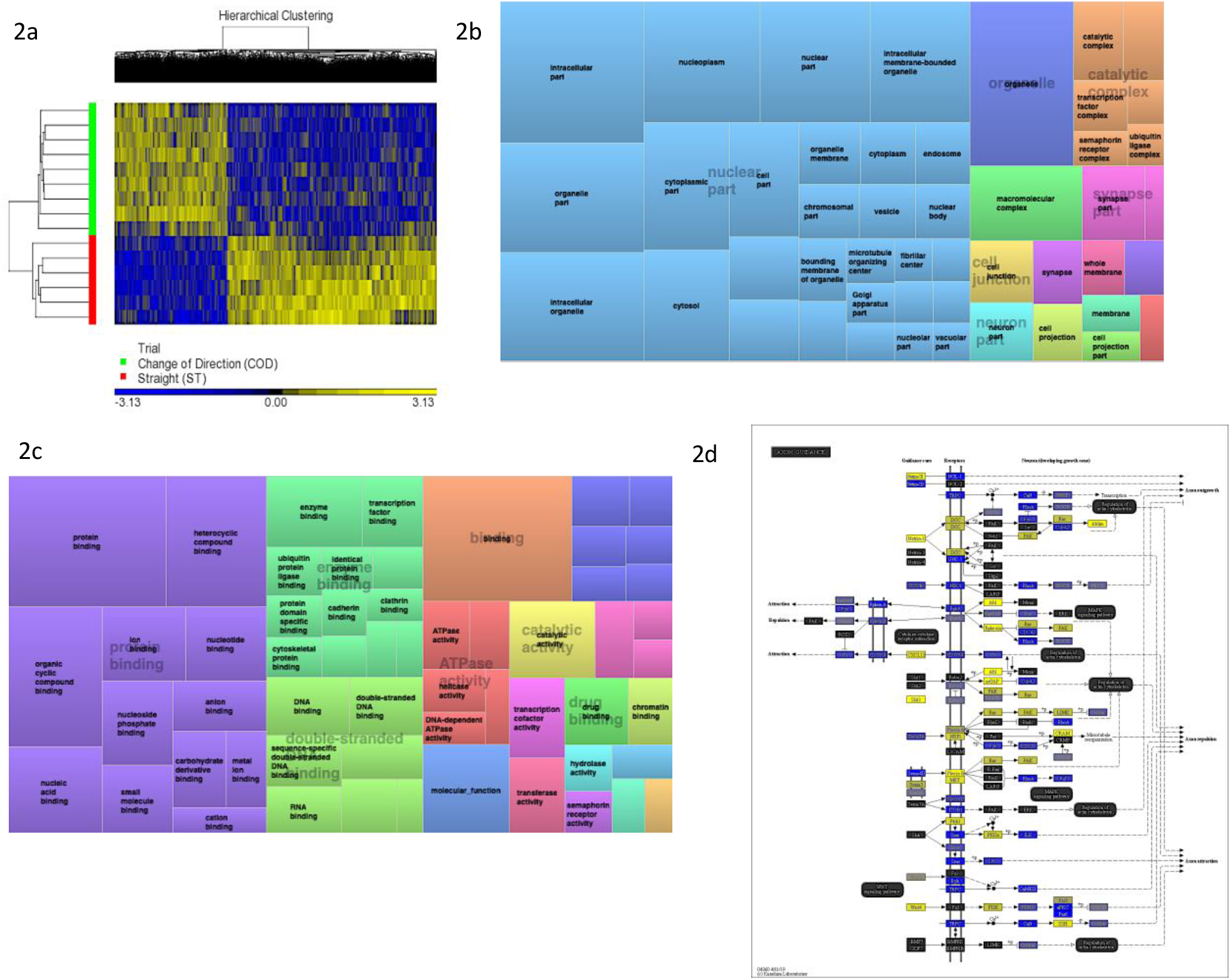
**a.** Hierarchical heat map for analysis of DMPs in COD vs. ST trials across all time. **b.** DMP enrichment TreeMap of overarching Gene Ontologies (GO) term ‘cellular component’ in COD vs. ST trials. Demonstrating top enriched GO terms of ‘intracellular part’, ‘organelle’ and ‘intracellular organelle part’. **c.** DMP enrichment TreeMap of overarching Gene Ontologies (GO) ‘molecular function’. Demonstrating top enriched GO terms of ‘protein binding’ and ‘binding’. **d.** Illustration of the top KEGG pathway ‘axon guidance’ demonstrating predominantly hypomethylation (blue) in COD vs. ST sprint trials. Note, the most significant (lowest p-value) DMP for each gene is used to colour this pathway image. Therefore, this is not always accurate where multiple DMPs occur for a single gene and the image is therefore only a visual representation of the overarching methylation profile in this pathway. Full and accurate DMP lists for the axon guidance pathway in these conditions can be found in (**Suppl. File 3i**).

### Change of direction exercise evokes the largest hypomethylated signature immediately post exercise compared with straight line exercise, particularly in protein binding, axon guidance and insulin related pathways

A 2-Way ANOVA for ‘Trial’ x ‘Time’ interaction, that demonstrates a difference in methylation between COD and ST sprint exercise immediately post and 24 hrs post exercise, identified **7,844** significant DMPs **(Suppl. File 4a**). 1,349 DMPs were promoter associated with 1,846 DMPs within CpG islands. Contrasts for COD post vs. ST post identified **6,489** significant DMPs (**Suppl. File. 4b**). Where hypomethylation was predominant post exercise in COD vs. ST post exercise, with 4,472 hypomethylated and 2,017 hypermethylated DMPs **(Figure 3a)**. At 24 hrs (COD 24 h vs. ST 24 h) there were **6,647** significant DMPs (**Suppl. File 4c**), where the shift in favouring hypomethylation post exercise was now more balanced with hypermethylation at 24 h, yet still with a larger total of hypomethylated (3,649) compared with hypermethylated (2,953) DMPs (**Figure 3b**). Furthermore, SOM analysis of mean temporal changes generated over time from the mean of each condition using the interaction Trial x Time (7,844 DMP) list demonstrated the following temporal profiles depicted in **Figure 3c** for COD exercise and **Figure 3d** for ST exercise. Indeed, these profiles confirmed that the majority of DMPs in the COD trial demonstrated a hypomethylated compared with hypermethylated profile particularly in the immediately post exercise timepoint.

**Figure 3.**
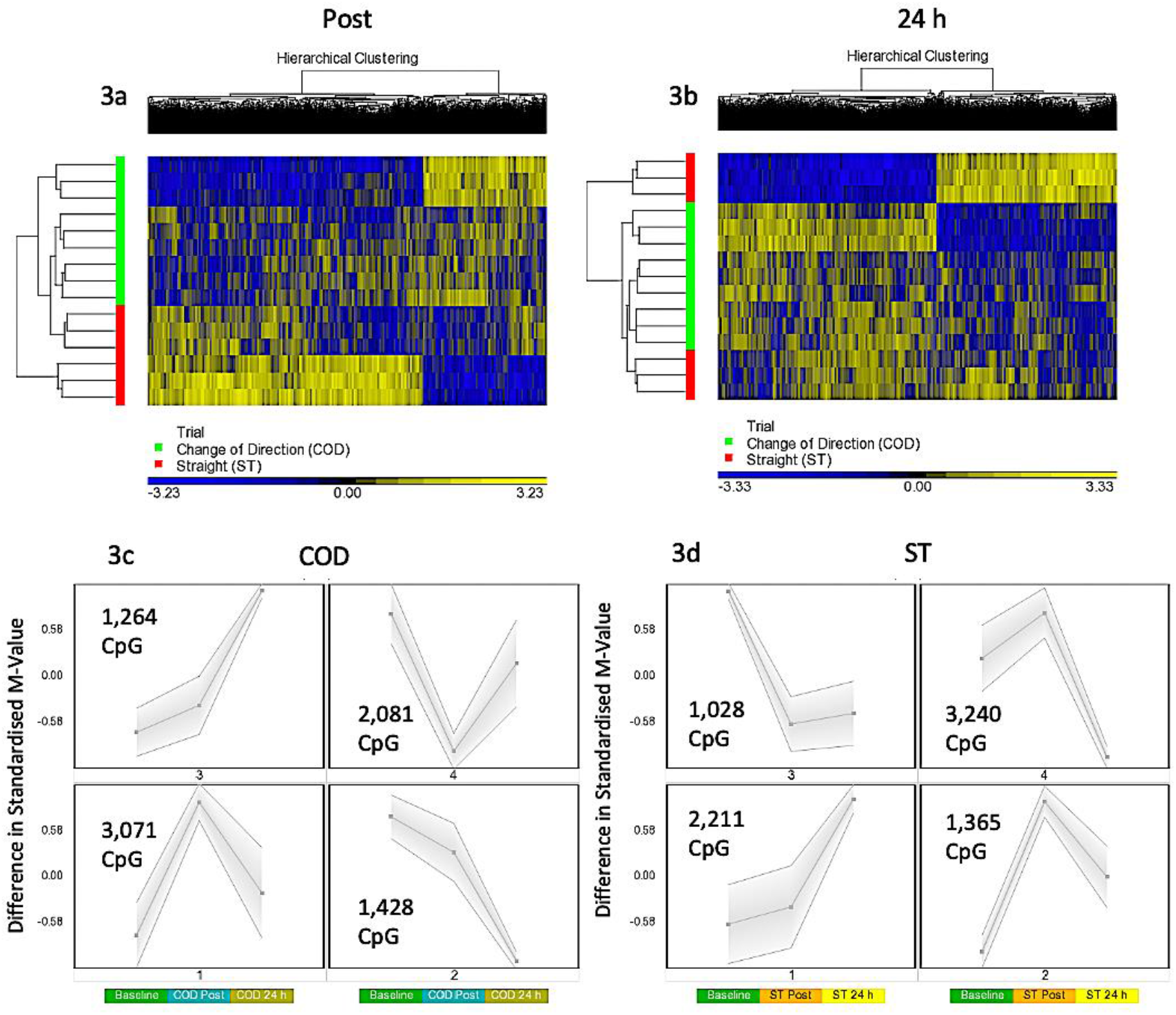
**a.** Hierarchical heat map for analysis of DMPs in COD vs. ST trials immediately post exercise. **b.** Hierarchical heat map for analysis of DMPs in COD vs. ST trials 24 h post exercise **c.** SOM temporal analysis of DMPs over exercise time in COD exercise trial at baseline, immediately post and 24 hr post exercise. **d.** SOM temporal analysis of DMPs over exercise time in ST exercise trial at baseline, immediately post and 24 hr post exercise.

In gene ontology (GO) analysis of the 2 Way ANOVA ‘Trial’ x ‘Time’ interaction (**7,844 DMP list; Suppl. File 4d**), identified overarching GO terms: Biological process, cellular component, and molecular function, as well as the top GO term protein binding (within molecular function) (**Suppl. File 4e**) that demonstrated more CpG’s hypomethylated compared with hypermethylated (2,276 hypomethylated versus 1,386 hypermethylated). The same trend was observed after KEGG pathway analysis (**Suppl. File 4f**) where the top differentially enriched pathway was ‘axon guidance’ (**Suppl. File 4g**) with more CpG’s hypomethylated in COD post vs. ST post exercise (61 CpG hypomethylated, 44 CpG hypermethylated).

More specifically post exercise in COD vs. ST trials, gene ontology (GO) analysis of the differentially methylated CpG’s within this contrast (**6,489 DMP list** above; **Suppl. File 4h**) demonstrated the most significantly enriched top 5 GO terms as: 1) Protein binding (**Suppl. File 4i**), 2) binding, 3) positive regulation of cellular metabolic process, 4) positive regulation of metabolic process, and 5) positive regulation of macromolecule metabolic process (**Suppl. Files 4j, k, l, m**) that all demonstrated more CpG’s that were hypomethylated vs. hypermethylated in these GO terms. By 24 hrs in the **6,647 DMP list** (contrast COD 24 h vs. ST 24 h) there was a shift back to a more balanced hypo/hypermethylated ratio in the top 5 GO terms (**Suppl. File 4n**) for: 1) protein binding (**Suppl. File 4o**), 2) binding, 3) regulation of signaling, 5) regulation of cell communication, and 5) anatomical structure morphogenesis; **Suppl. File 4p, q, r, s**).

The same trend was observed after KEGG pathway analysis (**Suppl. File 4t**) where the top differentially enriched hypomethylated pathways were: 1) Insulin resistance, 2) endocytosis, 3) axon guidance, 4) MAPK signalling pathway and, 5) insulin signaling pathway in the COD post vs. ST post contrast (**Suppl. Files 4u, v, w, x, y**). In the 24 h vs. 24 h ST trial contrast KEGG pathway analysis (**Suppl. File 5a**) with more balanced hypo/hypermethylation ratio: 1) ErbB signaling, 2) Non-small cell lung cancer, 3) axon guidance, 4) proteoglycans in cancer, and 5) glioma (**Suppl. Files 5b,c,d,e,f**). In DMR analysis COD post vs. ST post (**Suppl. File 5g**), identified enriched hypomethylation in genes: CIITA, PRR5, WDR46/PFDN6, MAGI2, RNF167 and DGKZ. In COD 24 h vs. ST 24 h DMR analysis suggested genes COL1A1 and CHID1 possessed enriched hypermethylation (**Suppl. File 5h**).

### Comparative methylome and transcriptome analyses identifies inverse relationships in DNA methylation and gene expression: NR4A1 is hypomethylated and associated with increased gene expression after acute exercise

In order to identify genes that were differentially methylated and also altered at the gene expression level after exercise between COD and ST sprint exercise trials, we first overlapped the present studies methylome data (significant DMPs main effect for ‘time’ **7,612 CpG list**, main effect for ‘trial’ **13,281 CpG list** and the interaction ‘time’ x ‘trial’ **7,844 CpG list**) with recent published transcriptome data sets following acute sprint exercise^13^. In this sprint transcriptome, the authors identified 879 genes that were significantly differentially expressed post-exercise (471 upregulated / 408 downregulated - List in **Suppl. File 6a**). Out of these 879 genes that had altered gene expression across the sprint transcriptome, there was large overlap of genes (49% or 431 genes) that were also differentially methylated in at least one of the significant DMP analyses in the present study (**Figure 4a**, **Suppl. File 6b** for gene list). To determine genes that were frequently occurring DMPs in COD vs. ST sprint exercise trials, that mapped through to significant differential gene expression in the transcriptome analysis, using Venn diagram analysis we first identified that out of these 431 genes, 61 were shared by all methylation analyses in the present study (**Figure 4a**, **Suppl. File 6c**). Eighteen of the genes that were upregulated, also displayed hypomethylation (on 27 DMPs) immediately post exercise in COD vs. ST sprint exercise condition (**Suppl. File 6d**). Fifteen of the genes, that were downregulated were also hypermethylated (24 DMPs) immediately post exercise in COD vs. ST conditions (**Suppl. File 6e**). Fifteen genes were also upregulated and hypomethylated (21 DMPs) 24 hrs post exercise (**Suppl. File 6f**) and twenty-one of these genes were downregulated and hypermethylated (30 DMPs) 24 hrs post exercise (**Suppl. File 6g**). Those genes with the same trend (down and hypermethylated) at both time points (post and 24hrs) in COD vs. ST exercise trials, included a final list of 13 genes (containing 28 DMPs) including: ATP2C1, CCDC88C, FAM111A, GAB1, HMCN1, LRRTM4, LTBP1, MAGI1, MYH10, MYO10, NEDD9, NIPAL3 and RHOBTB1 (**Suppl. File 6h**). With 10 genes (containing 22 DMPs) that were upregulated and hypomethylated at both post and 24 hrs in COD vs. ST sprint exercise trials, including genes: ADM, DUSP1, IRS2, MAP2K3, PDE2A, PDE4E, PLXNA2, POR, PPP1R15A and VEGFA (depicted **Figure 4b**, **Suppl. File 6i**). The majority of these hypomethylated and upregulated genes were associated with the MAPK pathway, together with the canonical angiogenesis gene, VEGFA (see additional DMR data for VEGFA below).

**Figure 4.**
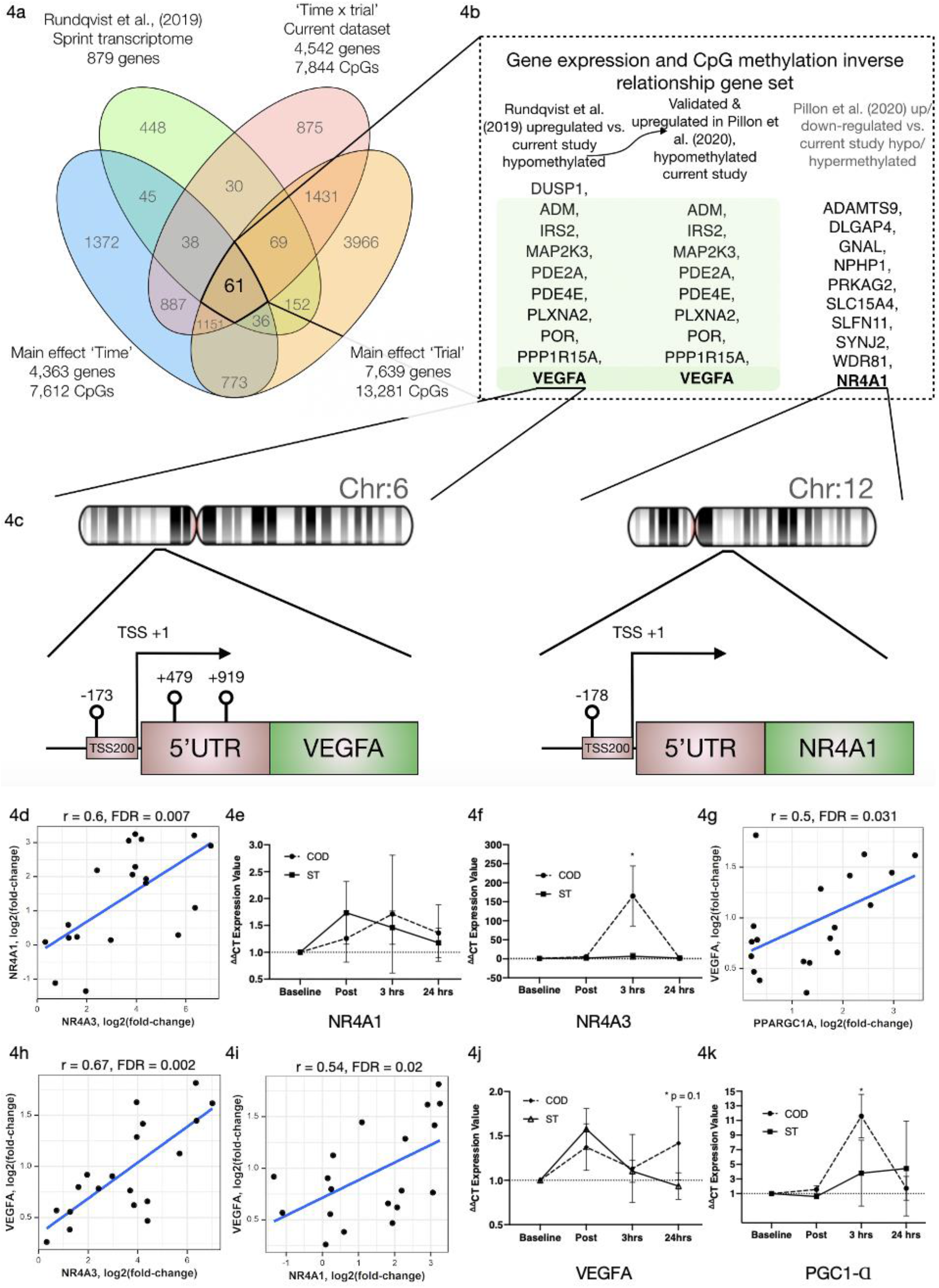
**a.** Venn diagram analyses of the overlap between the sprint transcriptome^13^ and differential methylome analysis in the present study. A total of 431 genes overlapped with the sprint transcriptome and were identified as significant DMPs in at least one of the methylome analyses in the present study. 61 genes were and up/down regulated in the sprint transcriptome and significantly differentially methylated across all methylome analyses in the present study. **b**. Genes (shaded in green) out of the 61 identified in figure 4a, that were significantly upregulated in the sprint transcriptome^13^, upregulated in acute aerobic exercise transcriptome meta-analysis^14^ as well as hypomethylated in all methylome analyses in the present study. Unshaded genes were upregulated in acute aerobic exercise transcriptome meta-analysis^14^ (but not changed in the sprint transcriptome) and significantly hypo or hyper methylated in all methylome analyses in the present study. **d.** Highly significant positive correlations in gene expression for NR4A1 vs. NR4A3 **e.** Gene expression of NR4A1 and **f.** NR4A3 at post, 3 hrs and 24 hrs after COD vs. ST exercise. Highly significant positive correlations in gene expression for **g**. VEGFA vs. PPARGC1A (PGC1-α), **h.** VEGFA vs. NR4A3 and **i.** VEGFA vs. NR4A1 in young healthy adult skeletal muscle immediately post up to 24 hours post exercise in new reanalysis of all acute aerobic transcriptomes meta-analyses^14^. Gene expression for **j.** VEGFA and **k.** PGC1-α at post, 3 hrs and 24 hrs after COD vs. ST exercise. * p ≤ 0.05 unless otherwise stated.

In addition, we looked to validate if the genes identified in the present study’s methylome and recent sprint transcriptome were also regulated in the most comprehensive set of acute aerobic transcriptomes to date^14^. Indeed, the vast majority of the 13 genes found to be hypermethylated/down-regulated and 10 genes hypomethylated/up-regulated in the present study’s methylome and sprint transcriptome^13^ were all significantly regulated (FDR < 0.05; Metamex.eu) at the gene expression level (except 2 genes; DUSP1 and NEDD9) in healthy skeletal muscle up to 24 hours after acute aerobic exercise in the transcriptome meta-analysis^14^. Most importantly, in the meta-analyses’ acute aerobic transcriptome data all these genes were regulated in same direction as identified above. This comparative methylome and transcriptome analysis therefore confirmed that there was an inverse relationship between these genes’ DNA methylation profile and their gene expression after acute exercise. Furthermore, re-analysing the full gene lists in the acute aerobic exercise transcriptomes meta-analysis data set^14^, we identified an additional 10 genes that were significantly up/down-regulated in the transcriptome meta-analyses (but not identified in the sprint transcriptome), that also demonstrated significantly altered methylation across all analyses in the present study (**Figure 4a**). These included: NR4A1, ADAMTS9, DLGAP4, GNAL, NPHP1, PRKAG2, SLC15A4, SLFN11, SYNJ2, WDR81 (**Figure 4b, Suppl. File 7a,b**). Notably, within this list, NR4A1 is a nuclear receptor and family member to NR4A3 that has been identified as one of the most exercise- and inactivity-responsive genes across all 66 published transcriptome data sets^14^. NR4A3 has a role in mediating the metabolic responses to exercise-like stimuli *in vitro*^14^. NR4A1 is also within KEGG pathway of ‘MAPK signalling’ and GO term ‘protein binding’ identified above as enriched hypomethylated pathways in COD vs. ST exercise. We identify that NR4A1, was hypomethylated in its promoter located in a CpG island close to its transcription start site TSS200 (cg11666140) in COD vs. ST exercise trials (**Figure 4c, Suppl. File 7a**), with another s-shelf associated CpG also identified (cg20661548) to be hypomethylated immediately post exercise in COD vs. ST trials (**Suppl. File 7b**). Further analysis shows that, alongside NR4A3, NR4A1 gene expression was also significantly upregulated (0.94 log fold) in exercise-transcriptome meta-analyses^14^ and within the top ten genes upregulated in the sprint transcriptome^13^. NR4A1 expression was also ranked 2^nd^ highest in its significant positive correlation with NR4A3 (r = 0.72, metamex.eu^14^) across the 66 published exercise and inactivity transcriptome data sets (metamex.eu^14^), with new analysis of NR4A1 and NR4A3 gene within meta-analyses transcriptomes of healthy young adult up to 24 hrs (to match the present study design) also suggesting a highly significant positive correlation (r = 0.6, FDR = 0.007; **Figure 4d**). Indeed, we were able to observe an elevation in NR4A1 at 3 hrs and 24 hrs post exercise in COD vs. ST trails, but this was not statistically significant (**Figure 4e**). Importantly however, we observed significant increases (p ≤ 0.05) of a large magnitude in NR4A3 gene expression (165.2 vs 6.8-fold, p ≤ 0.05) in COD vs. ST trial at 3 hours post exercise (**Figure 4f**). Therefore, the comparative analysis of our present methylome data and its overlap across recent sprint transcriptome^13^ and meta-analysis transcriptomes after acute aerobic exercise^14^, provides evidence to suggest that there is an inverse relationship between the above genes methylation and gene expression profile. Also identifying for the first time, that NR4A1 is an epigenetically regulated exercise gene and that NR4A3 is hugely upregulated (165-fold) as a consequence of performing high intensity change of direction compared with straight line sprint exercise.

### VEGFA has enriched hypomethylation in its promoter and is strongly associated with increased gene expression across exercise transcriptomes

We identified above that VEGFA was hypomethylated in the present study and gene expression increased across sprint and acute aerobic exercise meta-analysis transcriptomes. DMR analysis also identified enrichment of hypomethylation (3 DMPs) on the VEGFA gene (DMPs-cg01116220, cg04629501 and cg21099624- **Figure 4c, Suppl. File 8a**) in COD vs. ST exercise trails. All these DMPs were located amongst a CpG island within a 1093 base pair locus. Two of these DMPs were also within VEGFAs promoter region close to its transcription start site (TSS) and 1^st^ exon (cg21099624 - TSS200, cg01116220 - Exon 1, **Figure 4a**, **Suppl. File 8b**). Overall, suggesting that this gene has enriched hypomethylation in its promoter in COD compared with ST sprint exercise trials. Finally, as mentioned above, VEGFA expression was significantly increased in the sprint transcriptome data set^13^, but was also one of the most upregulated genes (0.96 log fold change; metamex.eu) in the transcriptome meta-analysis for acute aerobic exercise in healthy males up to 24 hrs after exercise^14^. VEGFA was also ranked top (1^st^) most significantly correlated gene at the expression level with the canonical exercise/metabolic regulators PGC1-α (r = 0.84, metamex.eu) and NR4A3 (r = 0.74, metamex.eu^14^) that was the most significantly upregulated across all modes of exercise^14^ and identified as hypomethylated in its promoter in the present study. Correlation analysis of VEGFA with PGC1-α and NR4A3 genes within meta-analyses transcriptomes of healthy young adults immediately post and up to 24 hrs (to match the current study design) post-acute aerobic exercise also suggested a significant correlation between VEGFA and PGC1-α (r = 0.5, FDR = 0.031; **Figure 4g**) and NR4A3 (r = 0.67, FDR = 0.002; **Figure 4h**). Further, VEGFA was significantly positively correlated with NR4A1 (r = 0.58, metamex.eu^14^) across all exercise transcriptomes and also after analysis of acute-aerobic exercise transcriptomes in healthy young adults (r = 0.54, FDR = 0.02; **Figure 4i**). Importantly, we identified VEGFA to possess enriched hypomethylated in its promoter in the current study. Interrogation of aged skeletal muscle tissue methylome in our recent studies, also suggests one of the same VEGFA CpG sites (cg04629501) is oppositely regulated (hypermethylated) with age^25^ compared with exercise (hypomethylated) in the present study. Therefore, VEGFA is hypomethylated in its promoter and is associated with its own increased gene expression (as well as highly correlated with PGC1-α, NR4A1 and NR4A3 expression) in the post-acute aerobic exercise transcriptome. Finally, VEGFA gene expression was elevated at 24 hours (p = 0.01) in COD vs. ST trials (**Figure 4J**). We also observed significant increases (p ≤ 0.05) of a large magnitude in canonical metabolic gene, PGC1-α (11.6 vs. 3.8-fold in COD vs. ST) in COD vs. ST trial at 3 hours post exercise (**Figure 4K**). Overall, for the first time this study identifies VEGFA as an epigenetically regulated gene in the response to acute exercise.

## Discussion

The present study aimed to investigate the methylome and transcriptome immediately post and 24 hrs after acute exercise in two sport and exercise relevant sprint shuttle running protocols: 1) Change of direction (COD) and, 2) straight line (ST) sprint exercise. We first demonstrated that while sprint trials were matched for classically defined intensity measures (speed x distance) and the number of acceleration and deceleration movements, by including changes of direction into the protocol elicited greater movement (GPS Playerload™), physiological (HR), metabolic (lactate), subjective central and peripheral (differential RPE) loading compared with straight line sprint running. Therefore, the comparison of COD versus ST sprint exercise represented a suitable model to investigate the time-course of the DNA methylome and comparative transcriptome following increased loading exercise in human skeletal muscle using physiologically relevant movements predominant within many sporting/exercise contexts and health prescription settings. Overall, we identified that both sprint exercise conditions evoked extensive alterations in the methylome immediately post and 24 hrs after exercise, particularly in MAPK, AMPK and axon guidance pathways. COD exercise evoked a greater hypomethylation response across the genome particularly enriched in: Protein binding, MAPK, AMPK, insulin and axon guidance pathways, specifically immediately post exercise, compared with ST exercise. Further, comparative transcriptome analysis with recent sprint running transcriptomes identified considerable 49% overlap of genes altered at the expression level, and that were also differentially methylated after COD exercise. In particular, after differential methylated region analysis of genes altered across the methylome, we identified that vascular endothelial growth factor A (VEGFA) and downstream nuclear transcription factor, NR4A1, possessed hypomethylation within their promoter regions. VEGFA and NR4A1 was also significantly upregulated in both sprint transcriptomes and recent comprehensive meta-analysis of 66 published exercise transcriptomes^14^. Furthermore, within these published meta-analyses, VEGFA was highest ranked gene positively correlated with well-established metabolic regulators; PCG1-alpha and NR4A3 expression in human skeletal muscle after exercise^14^. In a re-analysis of the meta-analysis transcriptome to correspond with our study design (acute exercise immediately up to 24 hrs post in healthy skeletal muscle), we also identified significantly positively correlations of VEGFA with PCG1-alpha and NR4A3 gene expression. We also confirmed increased gene expression of VEGFA at 24 hrs, PGC1-α and NR4A3 at 3 hrs post in COD exercise vs. ST exercise in the present study. In summary, we demonstrate that increased physiological load via change of direction sprint exercise in human skeletal muscle evokes considerable epigenetic modifications that are associated with changes in gene expression in genes responsible for adaptation to exercise. Further, incorporating changes in direction into exercise regimes overtime may ultimately help improve performance within a sporting context or health outcomes of populations with dysfunction in these gene pathways (MAPK, insulin signalling).

Acute COD sprint exercise evoked enriched hypomethylation in pathways such as insulin resistance and insulin signalling (specifically immediately post exercise) that have been demonstrated to be hypomethylated in skeletal muscle of people with type-II diabetes following 6 months of aerobic exercise training (3 days/wk),^10^ and in people with type-II diabetes that are non-responders to exercise^26^. This supports the notion that this type of change of direction sprint exercise maybe beneficial for those with metabolic disease, providing this type of higher intensity exercise is tolerable for these individuals. Others have suggested that it is well tolerated and also extremely practical, given a reduced time-burden of the exercise regime^27^. Furthermore, aging is associated with hypermethylation in skeletal muscle tissue^25, 28^ and muscle stem cells^25^. With increased physical activity and resistance exercise demonstrated to somewhat reverse the hypermethylated signature observed with age^25, 29^. Therefore, as COD exercise evokes greater hypomethylation than ST running, change of direction exercise may also have a beneficial epigenomic impact in reversing the hypermethylation observed in skeletal muscle tissue of aged individuals. Indeed, re-interrogation of the methylome in aged skeletal muscle tissues, reveals one of the same VEGFA CpG sites (cg04629501) is hypermethylated with aging^25^ as compared with the hypomethylated response seen with exercise in the present study. Something, that requires future investigation with COD exercise in elderly populations.

Despite COD exercise evoking a larger hypomethylation response versus straight line running. One of the surprising findings of the present study was the temporal exercise alone methylome response (regardless of experimental trial) demonstrated that there was a hypermethylation response immediately post exercise (30 minutes), yet after 24 h, a large majority of the same DMPs were hypomethylated, suggesting a switch in the methylation status of these DMPs. Future work could perhaps investigate changes in DNA methyltransferase activity at the protein level, such as DNMT3A and DNMT3B to investigate whether this ‘transient methylation’ phenomena is occurring. However, it is worth mentioning that the effect of the different trials is not included in the exercise alone analysis (i.e. a main effect for ‘time’), that highlighted this response. Therefore, the significant differences between the trials in terms of movement patterns, physiological and metabolic loads may contribute to this finding, and therefore warrants further investigation.

At the candidate gene level, in the present study we identified that Nuclear Receptor Subfamily 4 Group A Member 1 (NR4A1), also known as nerve growth factor IB (NGFIB) or Nur77, was hypomethylated in its promoter in COD vs. ST exercise trials. NR4A1 is also associated with GO ‘protein binding’ and KEGG pathway ‘MAPK signalling’ identified to have enriched hypomethylation at the pathway level in COD vs. ST trials. NR4A1 is member of the steroid-thyroid hormone-retinoid receptor superfamily, where the encoded protein acts as a nuclear transcription factor. NR4A1 has been shown to be increased in response to various signalling events including; cAMP, growth factors, mechanical stress, cAMP, calcium, and cytokines^30^. NR4A1 is the most abundant nuclear receptor in skeletal muscle as compared with NR4A2 and 3, and its expression is higher in fast-twitch muscle^31^. NR4A1 abundance increases with beta-adrenergic stimulation and after exercise^31, 32, 33, 34, 35^. NR4A1 is associated with glucose uptake, glycolysis and glycogenolysis and^31^ its global knock-out associated with an increased predisposition to obesity, insulin resistance and reduced muscle mass^36, 37^. Overexpression of NR4A1 in skeletal muscle increases oxidative metabolism^38^. Indeed, NR4A1 is also significantly upregulated (0.94 log fold) in exercise-transcriptome meta-analyses^14^ and within the top ten genes upregulated in the sprint transcriptome^13^. NR4A1 expression was also ranked 2^nd^ highest in its significant positive correlation with its other family member NR4A3 (r = 0.72) in a meta-analysis of exercise transcriptome data sets (metamex.eu^14^). Indeed, NR4A3 was the most significantly altered gene across all meta-analyses data sets and has been confirmed to be responsive to exercise stimuli in-vitro^14^. In the present study we also confirmed a robust 165-fold increase in NR4A3 in COD vs. ST trail at the gene expression level. Finally, NR4A1 methylation has also been linked to epigenetic memory of skeletal muscle in the offspring of high-fat fed mothers and that this memory could be reversed if the offspring undertook exercise^39^. Given that hypomethylation can be retained after acute resistance exercise and training^2^, and given the association of NR4A1 with an epigenetic memory of nutrient stress suggests that the role of NR4A1 in an epigenetic memory of aerobic exercise in adult human skeletal muscle requires future investigation.

We demonstrate that Vascular Endothelial Growth Factor (VEGF), a potent angiogenic factor originally described in vascular endothelial cells (also expressed in skeletal muscle), has enriched hypomethylation in its promoter close to its transcription start site in change of direction versus straight line sprint exercise. VEGF is involved in training-induced capillary growth^40, 41, 42^ and is increased after acute exercise^43, 44, 45^ with protein levels elevated during the first 4 weeks of exercise training^46, 47^. However, levels are lower in muscle of elderly individuals, but enhanced with training^48, 49^. In the present study we also confirmed a larger increase in VEGFA gene expression at 24 hrs in COD vs. ST trials. VEGFA was also ranked top (1^st^) most significantly correlated gene at the expression level with the canonical exercise/metabolic regulators PGC1-α (r = 0.84) and NR4A3 (r = 0.74) that was the most significantly upregulated across all modes of exercise in Pillion et al., (2020) across 66 published exercise transcriptome data sets. Re-analysis of the meta-analysis transcriptome to match our study design (acute exercise immediately up to 24 hrs post in healthy skeletal muscle), we also identified significantly positively correlations of VEGFA with PCG1-alpha and NR4A3 gene expression. We also demonstrate an increase, of a very large magnitude in PGC1-α (11.6 vs. 3.8-fold), as well as in NR4A3 (165.2 vs 6.8-fold) in COD vs. ST trials in the present study. VEGF mediates the upregulation of NR4A1 via activation of the PKD/HDAC7/MEF2 pathway^50, 51^ and addition of VEGF to endothelial cells increases NR4A1, 30-fold^52^. Therefore, to the best of our knowledge, this is the first study to demonstrate that both VEGFA and NR4A1 are epigenetically modified (hypomethylated) and associated with an increase in gene expression after acute exercise. This hypomethylation effect is enhanced in VEGF specifically after COD exercise compared with ST exercise. Our data raises the possibility that repeated COD exercise may improve VEGFs regulation of capillary formation in response to exercise. Therefore, future studies should undertake training with COD versus ST exercise in order to confirm whether the enhanced epigenetic change in VEGF leads to improved capillary formation.

It is important to note while there was a smaller cohort used in the present study for the discovery of differentially methylated sites, the significant overlap of this methylome analysis with recent sprint transcriptome datasets and across the transcriptome meta-analyses across 66 published studies, as well the identification of an inverse relationship in methylation and gene expression, significantly strengthens the findings and the potential applicability of the results to larger cohorts.

## Conclusion

We provide evidence that change of direction sprint exercise preferentially hypomethylates the skeletal muscle methylome as compared with straight line sprint exercise. Specifically, we found hypomethylation and increased gene expression of metabolic and angiogenic genes and pathways. The implication of this data suggests that introducing change of direction into high intensity running protocols could serve as an important modulator of a favourable epigenomic and transcriptomic landscape in response to exercise in athletes and trigger greater skeletal muscle remodelling through enhanced gene expression.

## Acknowledgments and funding

Mohd Firdaus Maasar received a funded PhD scholarship via the Malaysian government agency: Majlis Amanah Rakyat (MARA) via Barry Drust, Adam P. Sharples and Andrew Hulton. This work was supported by the UK’s Engineering and Physical Sciences Research Council (EPSRC) and the UK Medical Research Council’s (MRC) centre for doctoral training, via a studentship awarded Piotr Gorski in the group of Adam P. Sharples (PI). These data were also supported by a North Staffordshire Medical Institute (NMSI), the Society for Endocrinology and Glaxosmithkline grants awarded to Adam P. Sharples (PI). Funds from Liverpool John Moores University (UK) The Norwegian School of Sport Sciences, Oslo (Norway) and The University of Birmingham (UK) supported the PhD work by Mohd Firdaus Maasar, Daniel Turner, Piotr Gorski and Robert Seaborne in the groups of Adam P. Sharples and Barry Drust. Juleen R Zierath was supported from the Swedish Research Council for Sport Science (P2018-0097) and the Swedish Research Council (Vetenskapsrådet) (2015-00165). Nicolas J. Pillon was supported by an Individual Fellowship from the Marie Skłodowska-Curie Actions (European Commission, 704978).

## Declaration

All authors declare no conflicts of interest.

---

**Suppl. Figure 1.**
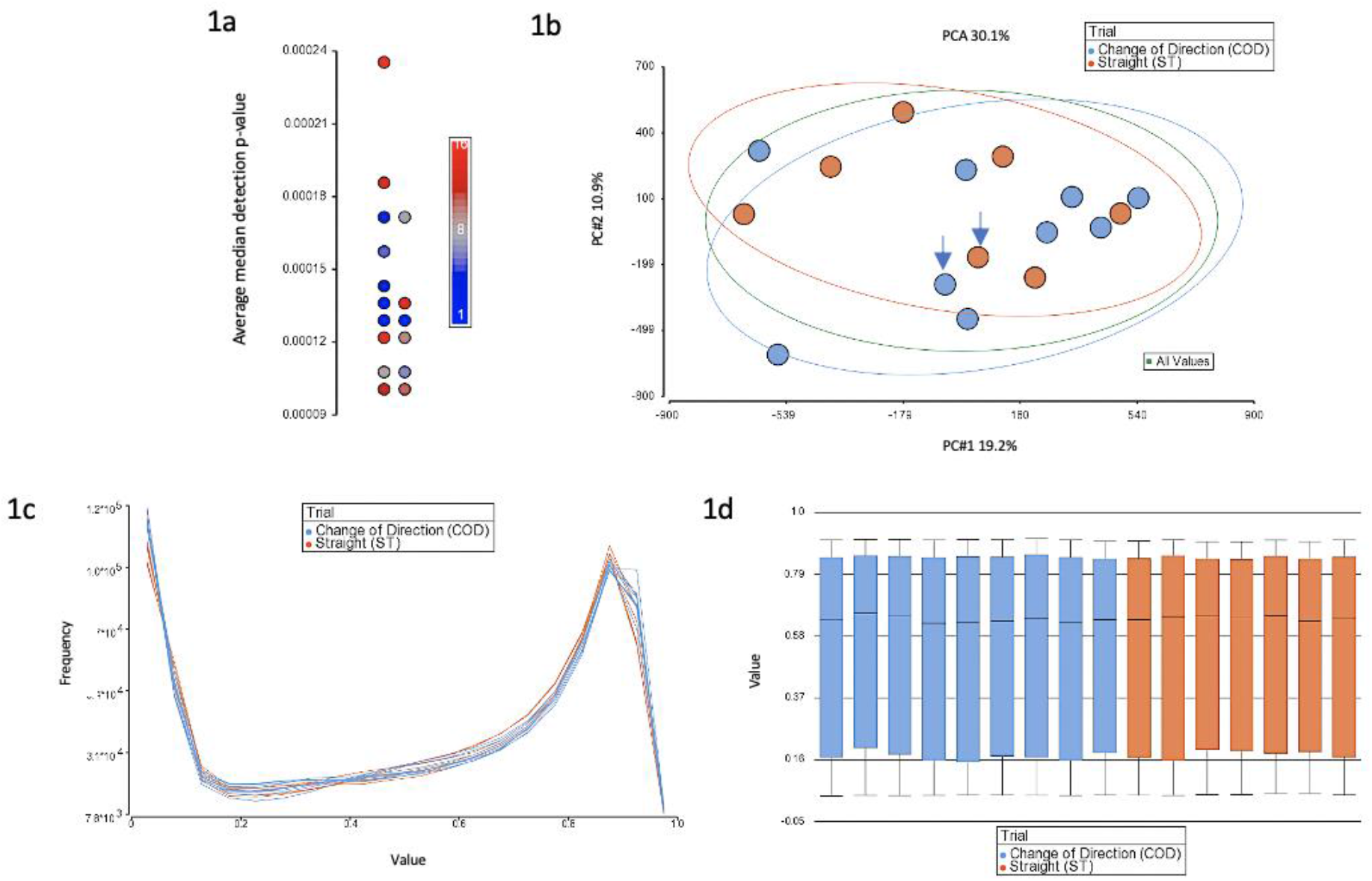

